# Intracellular mass density increase is accompanying but not sufficient for stiffening and growth arrest of yeast cells

**DOI:** 10.1101/388876

**Authors:** Shada Abuhattum, Kyoohyun Kim, Titus M. Franzmann, Anne Eßlinger, Daniel Midtvedt, Raimund Schlüßler, Stephanie Möllmert, Hui-Shun Kuan, Simon Alberti, Vasily Zaburdaev, Jochen Guck

**Affiliations:** Biotechnology Center, Center for Molecular and Cellular Bioengineering, Technische Universität Dresden, Dresden, Germany; JPK Instruments AG, Berlin, Germany; Max Planck Institute of Molecular Cell Biology and Genetics, Dresden, Germany; Department of Applied Physics, Chalmers University of Technology, Gothenburg, Sweden; Max Planck Institute for the Physics of Complex Systems, Dresden, Germany

**Keywords:** Yeast, Optical diffraction tomography, AFM, Refractive Index, Stiffness, Liquid solid transition.

## Abstract

Many organisms, including yeast cells, bacteria, nematodes and tardigrades, endure harsh environmental conditions, such as nutrient scarcity, or lack of water and energy for a remarkably long time. The rescue programs that these organisms launch upon encountering these adverse conditions include reprogramming their metabolism in order to enter a quiescent or dormant state in a controlled fashion. Reprogramming coincides with changes in the macromolecular architecture and changes in the physical and mechanical properties of the cells. However, the cellular mechanisms underlying the physical-mechanical changes remain enigmatic. Here, we induce metabolic arrest of yeast cells by lowering their intracellular pH. We then determine the differences in the intracellular mass density and stiffness of active and metabolically arrested cells using optical diffraction tomography and atomic force microscopy. We show that an increased intracellular mass density is associated with an increase in stiffness when the growth of yeast is arrested. However, increasing the intracellular mass density alone is not sufficient for maintenance of the growth-arrested state in yeast cells. Our data suggest that the cytoplasm of metabolically arrested yeast displays characteristics of solid. Our findings constitute a bridge between the mechanical behavior of the cytoplasm and the physical and chemical mechanisms of metabolically arrested cells with the ultimate aim of understanding dormant organisms.

## Introduction

Eukaryotic cells form compartments and organelles to spatially and temporally organize their cellular contents [1,2]. Many of these compartments such as nuclei, lysosomes, mitochondria and endoplasmic reticulum are bound by a lipid bilayer. Yet, recent studies have suggested that several types of organelles are not delimited by a lipid bilayer, but formed by liquid-liquid phase separation, where the dense phase of macromolecules such as RNAs and proteins are concentrated spontaneously from the surrounding solution [3,4]. A growing number of organelles have been reported as such membrane-less compartments, including nucleolus for ribosome biogenesis and stress response, and P bodies for RNA storage and degradation [2]. The liquid-liquid phase separation process can be tightly regulated by the cell owing its sensitive dependence on internal material concentration. However, under certain circumstances, these membrane-less compartments can also undergo liquid-to-solid phase transitions to form aberrant solid aggregates. For instance, when the genes encoding for the constituting macromolecules carry specific mutations, this phase transition is associated with neurodegenerative diseases via amyloid-like assembly of RNA binding protein Fused in Sarcoma (FUS) [5].

Liquid-to-solid phase transitions are not always pathological. Some organisms change their physical properties from a liquid- to solid-like state when faced with unfavorable conditions to facilitate survival. In this state, typically referred to as dormancy, the cell cycle is arrested and metabolic activity is reduced or suspended, but the cell can still recover to normal physiology and function when the conditions are favorable again. For example, plants develop seeds that survive cold and dry weather and germinate in spring [6]. Bacteria form metabolically inactive spores that are highly resistant to stress and antibiotics [7]. *Caenorhabditis elegans* enter a dauer stage that relies entirely on internal energy sources for as long as 4 months [8]. And finally, yeast cells compact their cytoplasm and cease proliferation in the absence of energy [9,10]. These changes in the function of the organism are associated with drastic alterations of the cellular architecture.

Munder et al. showed recently that lowering the intracellular pH of yeast cells, which is a consequence of lack of energy, promotes entry into dormancy [11]. When the cytosol was in acidic pH, the cells reduced their volume and formed macromolecular assemblies. These cells exhibited a reduction in the mobility of organelles and exogenous tracer particles, and were measurably stiffer than active cells. Yet, it is still unclear how they acquire this increased mechanical stability associated with dormancy. Is an increase in the intracellular mass density sufficient for a liquid-to-solid transition, which could then be associated with a glass-transition above a critical volume fraction? Or does it involve the assembly of macromolecules to form a percolated, solid-like intracellular matrix? In this study, we adopt the same technique of Munder et al. to arrest growth in fission yeast cells by lowering intracellular pH. We evaluate the physical properties of yeast cells optical diffraction tomography (ODT) and atomic force microscope (AFM) for probing their mass density and mechanical properties, respectively. We show that metabolically arrested cells exhibit higher intracellular mass density as well as elevated cell stiffness. However, an increased intracellular mass density alone is not sufficient to maintain cells in growth arrest. Thus, we suggest that yeast, when challenged by extreme conditions, utilizes distinct supramolecular structures in its dense cytoplasm that induce an increased mechanical stability.

## Materials and methods

### Yeast cell culture and growth assay

*S. pombe* strain ED668 was grown and maintained on YES (Formedium) agar plates at low temperatures. Two days prior to measurement, 50 mL liquid YES and 50 mL YES supplemented with 1.2 M sorbitol (Sigma) were inoculated with single colonies and incubated shaking at 180 rpm at 30°C overnight, respectively. The overnight cultures were then used to inoculate both, 50 mL YES and 50 mL YES supplemented with 1.2 M sorbitol at an optical density of OD_600 nm_ ~ 0.05, respectively and grown to early mid-log phase OD_600 nm_ ~ 0.2-0.4. The later was intended for adapting the cells to a hypertonic growth medium. Note that YES is a rich medium that contains glucose.

### pH adjustment of cells

The pH adjustment of the yeast cytosol was carried out as described previously [10,11]. Briefly, a set of either 100 mM potassium phosphate, 2% glucose, or 40 mM PIPES, 2% glucose was prepared at different pH values. Yeast cells from a liquid culture were centrifuged for 10 s at 2000 rcf and washed once with the buffer to remove residuals of culture medium. The cells were then resuspended in the respective buffer supplemented with 2 mM 2,4-dinitrophenol (DNP) in order to equilibrate the intracellular pH with the extracellular pH. Omitting DNP from the sample served as controls. The pH values used range between 6.0 and 7.6.

### Spheroplasting (cell wall removal)

Early mid-log cells were harvested by centrifugation (4 minutes at 1,500 rcf) and resuspended in phosphate buffer supplemented with 1.2 M sorbitol and 2% glucose. To remove the cell wall, 0.5 mg/mL Zymolyase 100T (ZymoLabs) and 2.5 mg/mL lysing enzymes from Trichoderma harzianum (Sigma) were added to the PBS medium and the reaction was incubated for approximately 120 minutes. Spheroplasts were then resuspended in 100 mM potassium phosphate buffer pH 6.0 or pH 7.5, respectively. 2 mM DNP were added as described in the previous section. Spheroplasting removes the very stiff cell wall (but not the cell membrane) of yeast cells so that the cell stiffness can be determined by indentation tests. Sorbitol is required in the medium after spheroplasting to prevent the cells from bursting.

### Optical diffraction tomography

The three-dimensional (3D) refractive index (RI) distribution of samples was measured using a custom-made optical diffraction tomography microscope. The optical setup of ODT employs Mach-Zehnder interferometry in order to measure complex optical fields of light scattered by samples from various incident angles, as shown in (Fig. 1A). A coherent laser beam (*λ* = 532 nm, frequency-doubled Nd-YAG laser, Torus, Laser Quantum, Inc., UK) is divided into two beams by a 2 × 2 single-mode fiber optic coupler. One beam is used as a reference beam and the other beam illuminates the specimen on the stage of a custom-made inverted microscope through a tube lens (*f* = 175 mm) and a high numerical aperture (NA) objective lens (NA = 1.2, 63×, water immersion, Carl Zeiss AG, Germany). To reconstruct a 3D RI tomogram of samples in a field-of-view, the samples are illuminated with 150 various incident angles scanned by a dual-axis galvanomirror (GVS012/M, Thorlabs Inc., USA). The diffracted beam from a sample is collected by a high NA objective lens (NA = 1.3, 100×, oil immersion, Carl Zeiss AG) and a tube lens (*f* = 200 mm). The total magnification is set to be 90.5×. The beam diffracted by the sample interferes with the reference beam at an image plane, and generates a spatially modulated hologram. The hologram is recorded with a CCD camera (FL3-U3-13Y3M-C, FLIR Systems, Inc., USA).

**Figure 1.**
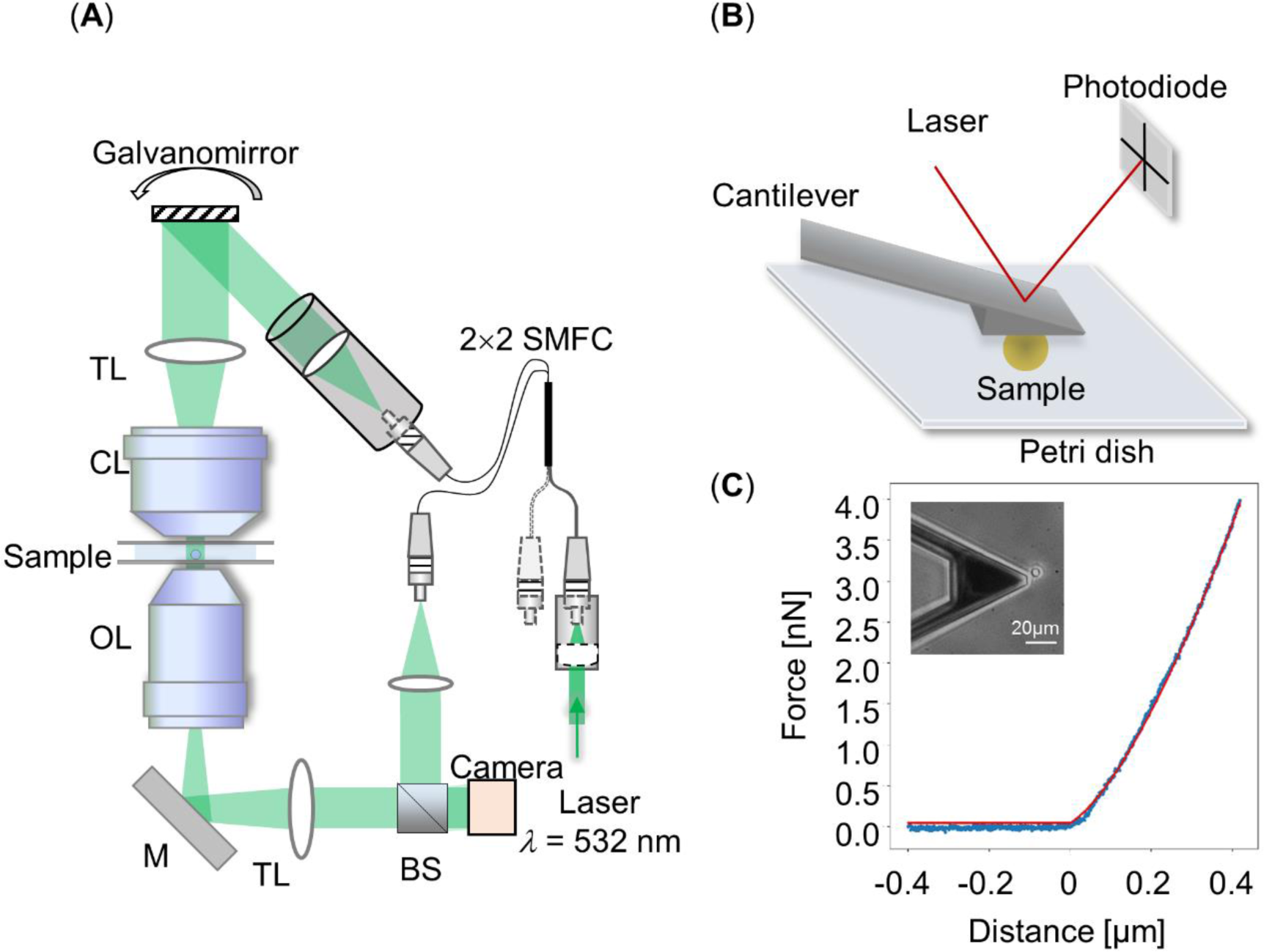
Experimental setups. **(A)** Schematic diagram of the optical setup for optical diffraction tomography (ODT). SMFC: single-mode fiber coupler, TL: tube lens, CL: condenser lens, OL: objective lens, M: mirror, BS: beam splitter. **(B)** Schematic diagram of atomic force microscopy (AFM) using a wedged cantilever to indent a spherical sample. **(C)** Representative force-distance curve (blue dotted line) recorded during compression with a wedged cantilever shown in inset. Hertz model fit (red solid line) to extract the apparent Young’s Modulus.

The complex optical fields of light scattered by the samples were retrieved from the recorded holograms by applying a Fourier transform-based field retrieval algorithm as previously published [12,13]. The 3D RI distribution of the samples was reconstructed from the retrieved complex optical fields via the Fourier diffraction theorem. A more detailed description of tomogram reconstruction can be found elsewhere [14–17]. From the reconstructed RI tomograms, the individual yeast cells were segmented by applying Otsu’s method and a watershed segmentation algorithm, and the mean RI value inside the binary image of an individual yeast cell was determined. The mass density of individual yeast cells is directly calculated from the mean RI value, since the RI value in biological samples is linearly proportional to the mass density inside cells as *n*(*x*,*y*,*z*) = *n*_m_ + *α*C(*x*,*y*,*z*), where *n*(*x*,*y*,*z*) is the 3D RI distribution of samples, *n*^m^ is the RI value of the surrounding medium (*n*_m_ = 1.339 at *λ* = 532 nm), *α* is an RI increment (*α* = 0.190 mL/g for protein [18]), and C(*x*,*y*,*z*) is the mass density inside cells. All tomogram acquisition and data analysis were performed using a custom-written MATLAB code.

### Atomic force microscopy (AFM) measurements

AFM-based nanoindentation measurements were performed using a Nanowizard 4 (JPK Instruments) and an optical inverted microscope (Observer D1, Zeiss). For immobilizing the spheroplasts of *S. pombe* prior to the measurement, CellTak (Corning), a cell adhesive protein solution, was first applied to the plastic petri dish and then rinsed for removing residuals [19]. Measurements were carried out using a wedged cantilever that is parallel to the petri dish as illustrated in (Fig. 1B). This simplifies indenting spherical cells, prevents their movement during the procedure and compensates for the original 10° angle of the glass blocks used for mounting the cantilevers. The wedged cantilever was prepared using a UV curing glue that was applied to tipless cantilevers (PNP-TR-TL, nominal spring constant *k* = 0.32 N/m, Nanoworld, or HQ: CSC37/tipless/No Al, nominal spring constant *k* = 0.3 N/m, Mikromasch). For determining cell stiffness, the cantilever was positioned above a single spheroplast and lowered with a speed of 5 µm/s. Force-distance curves were recorded until an indentation of 0.3 − 0.8 µm was reached (1.5 − 4 nN). All measurements were carried out in the phosphate buffer containing 1.2 M sorbitol, 2% glucose and DNP at 30°C.

### AFM-based indentation data analysis

The force-distance curves were analyzed using JPK data processing software. First, the curves’ baseline was adjusted and the data were corrected for the tip-sample separation [20]. Then each curve was fitted with the Hertz model modified by Sneddon for a spherical geometry to evaluate the spherical spheroplasts as shown in (Fig. 1C) [21,22] using

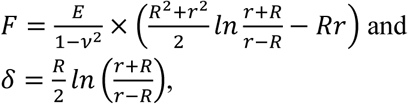

where *F* denotes the indentation force, *δ* the indentation depth, *r* the cell radius, *R* the radius of the circular contact area and ν is the Possion’s ratio and is set to 0.5 in all measurements.

The radius of the cells was measured for each of the cell groups and the average radius was used for each individual group (ranging between *r* = 2.28 − 2.45 µm) shown in the inset in Fig. 1C. The apparent Young’s modulus extracted from the fit was corrected for the additional stress that the cells encounter from the bottom of the plate, which causes an additional deformation of the spherical shape [23,24]. The Young’s modulus is reported as “apparent”, acknowledging the fact that some of the Hertz model assumptions are not met such as isotropic and homogeneous samples. While the absolute values are debatable, this approach still serves well the aim of quantitative comparison.

### Osmolality

Osmolality measurements of mediums and buffers used were carried out using an Osmomat 3000 Freezing point osmometer (Gonotec).Three independent measurements were carried out per sample and averaged to calculate the mean and standard deviation.

### Statistical analysis

To perform statistical analysis on ODT and AFM experiments we measured three independent sets of yeast cells. The mean values were subjected to statistical analysis by the two-tailed Mann-Whitney U-test using python scipy library. The shown asterisks indicate significance levels: ∗ *p* < 0.01, ** *p* < 0.001, *** *p* < 0.0001.

## Results

Yeast cells usually grow in acidic environments and use proton-translocating ATPases for maintaining their intracellular pH neutral at the expense of large amounts of energy in the form of ATP hydrolysis. When energy is scarce, however, these cells enter into a dormant state [25–28]. In dormancy, the cytosol becomes acidic as the supply of ATP is not sufficient to maintain the proton pumping [9–11]. We have shown that lowering the intracellular pH, by incubating the cells in acidic buffers and adding protonophores, such as DNP, that carry protons across the membrane, promotes entry into a dormant state in both fission and budding yeast cells, even if energy is present [11]. As a consequence of reducing intracellular pH, two main changes occur in the cell cytoplasm. First, a large fraction of cytosolic proteins reduces the net charge, becomes less soluble and forms aggregates, and second, the cell displays a reduction in volume and an increase in cytoplasmic crowding. Both processes can potentially lead to an increase in the mechanical stability of the cytoplasm in the liquid-to-solid transition that is observed, either mediated by the formation of a percolated cellular matrix of supramolecular assemblies (similar to a sol-gel transition) or by the increase in mass density after volume loss above a critical value, leading to a glass-like state. Here we focus on fission yeast and decouple these two scenarios by designing a set of experiments to independently change intracellular pH and volume of yeast cells. We start by examining the effects of either pH changes or osmotic pressure on mass density using ODT. We then combine both effects to understand their mutual contribution to growth arrest of cells. Finally, we compare the mass density to the stiffness measurements done using AFM to determine whether an increased mass density is sufficient to drive cells into the mechanically stable state associated with dormancy.

### Lowering intracellular pH increases mass density of yeast cells

We have previously shown that lowering the intracellular pH promotes entry into dormancy and is coupled with volume decrease [11]. However, it is still unclear whether cells with low intracellular pH simply expel water and increase overall intracellular mass density or whether they also decrease the non-aqueous cytoplasmic content. Here we investigated the effect of this pH reduction on intracellular mass density. We used ODT to measure the RI values from which we calculated directly the mass density (see Methods and materials section) of yeast cells in media with different pH values starting from neutral pH 7.6 to acidification of the cytoplasm at pH 6.0 (see Fig. 2A). The media contained DNP in order to equilibrate the intracellular pH with the extracellular pH. As a control, we incubated the cells at two pH values (pH 6.0 and pH 7.6) in the absence of DNP so that the intracellular pH in the cells remained neutral. Figure 2B shows that the mean RI values of yeast were increased in the pH range of 6.2 to 6.4 compared to the other, more neutral pH values. The sigmoidal fit illustrates the cooperativity of this behavior. In contrast, the dry mass stayed constant for all values of intracellular pH (see Fig. 2C), which suggests that lowering intracellular pH increases the intracellular mass density as a result of volume decrease (See Supplementary Figure 1). However, it still remained unknown whether a reduction in volume and the associated increase in intracellular mass density alone would lead to stiffness changes associated with cells entering dormancy.

**Figure 2.**
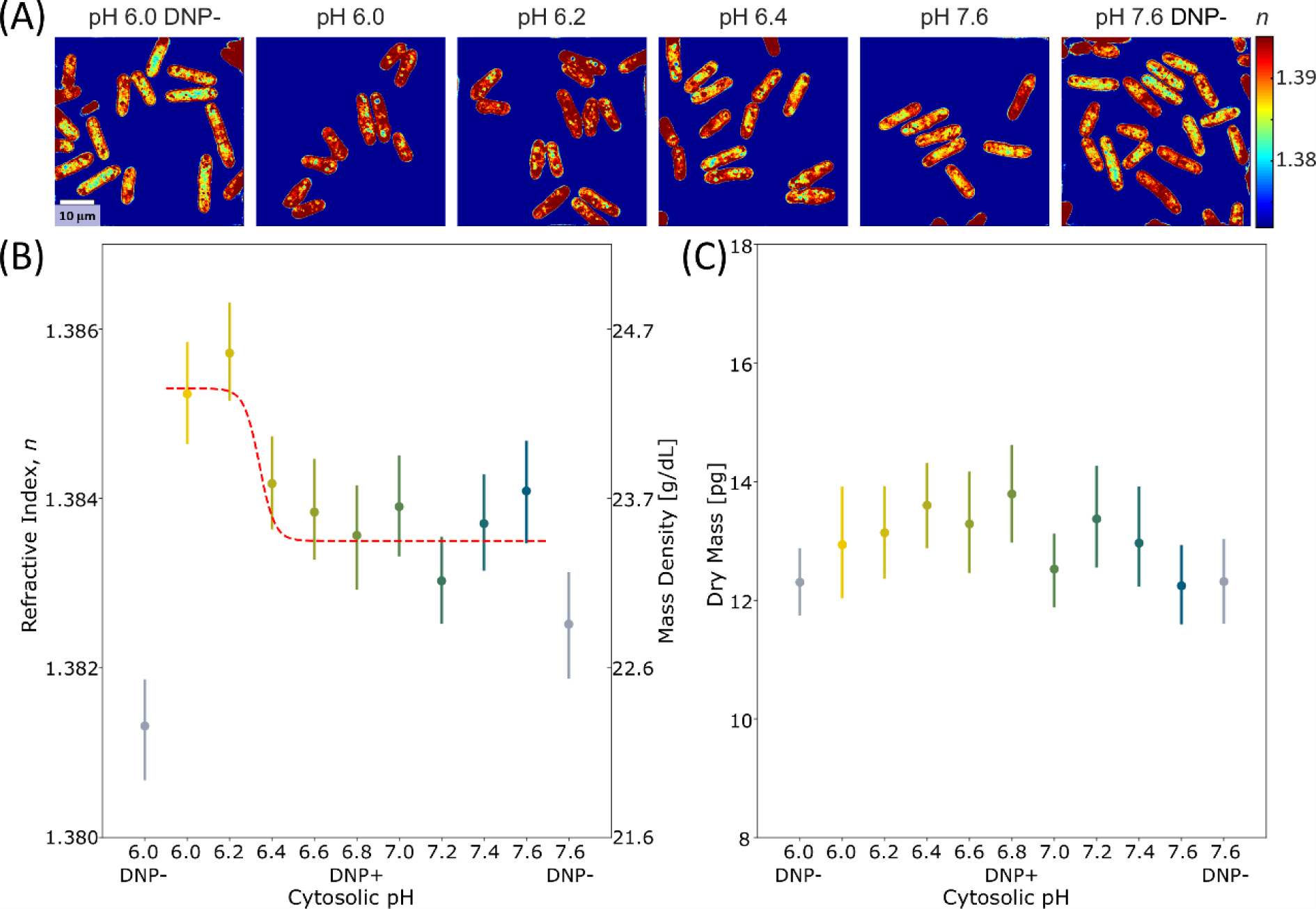
The effect of intracellular pH on the refractive index and dry mass of *S. pombe*. **(A)** Typical cross-sectional slices of the 3D refractive index tomograms of yeast cells in the *x*-*y* plane. Two control groups of cells incubated at pH 6.0 and 7.6 in the absence of DNP are denoted by DNP–. The scale bar indicates 10 μm. **(B-C)** The mean refractive index (RI) distribution **(B)** and the dry mass content **(C)** of yeast cells at different intracellular pH. the points shown correspond to the mean refractive index or dry mass measured at that pH value and lines indicate error bars (for each pH condition, *N* > 95).

### Cytoplasmic crowding is insufficient for promoting growth arrest

Next, we applied osmotic pressure on yeast cells without changing their intracellular pH to examine whether high intracellular mass density triggers dormancy. We measured RI tomograms of yeast cells grown in YES medium as a control (osmolality 196±1 mOsmol/kg) (see Fig. 3A), and compared their RI tomograms with those obtained for cells after adding 1.2 M sorbitol to YES medium (osmolality 1,946±8 mOsmol/kg) (see Fig. 3B) as well as with those for cells that had adapted to osmotic pressure overnight (Fig. 3C).

**Figure 3.**
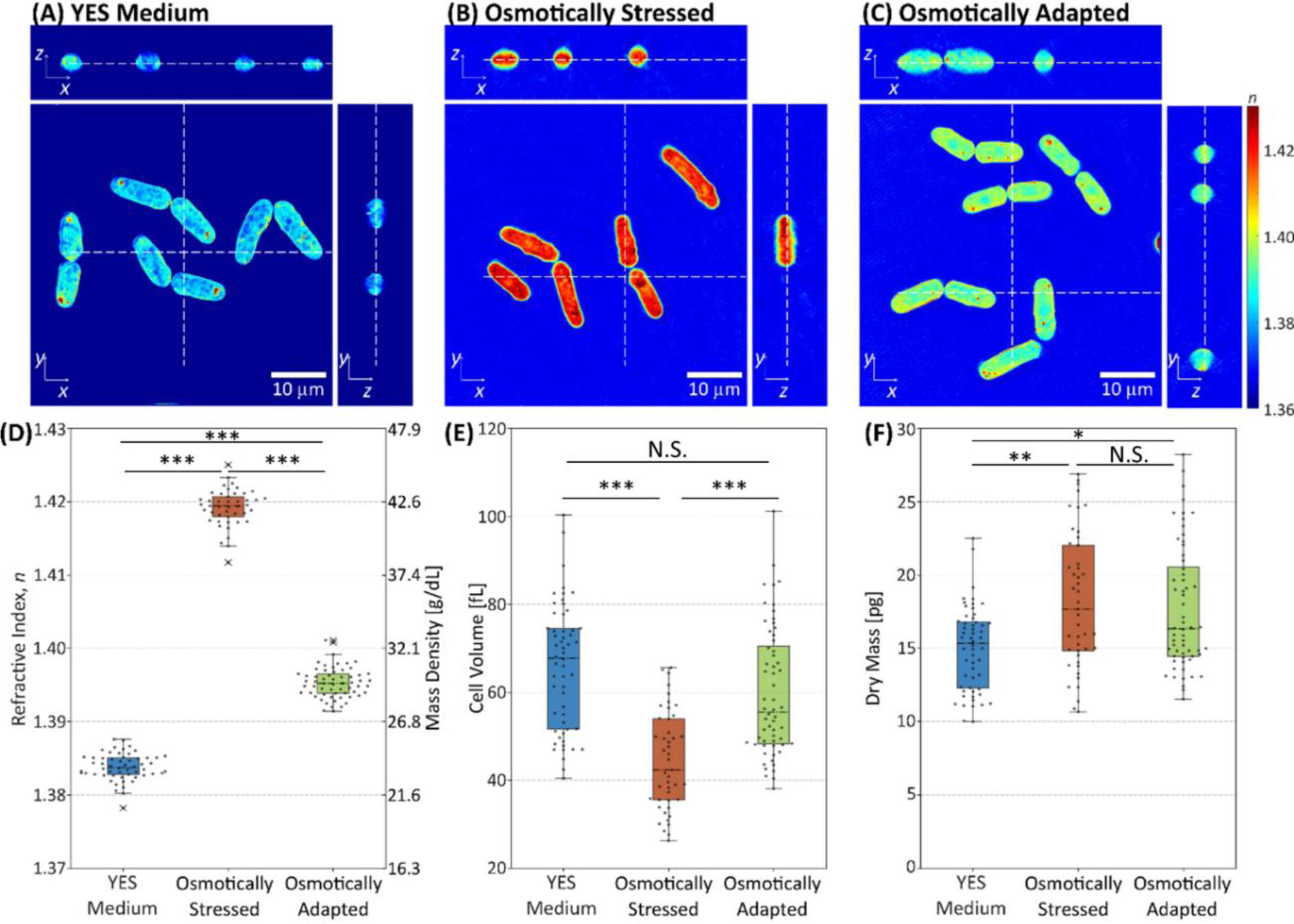
The effect of osmotic stress and adaptation on yeast cells. **(A-C)** Cross-sectional slices in the *x*-*y*, *y*-*z*, and *x*-*z* planes of 3D RI tomograms of **(A)** control in YES medium, **(B)** osmotically stressed, and **(C)** osmotically adapted yeast cells. The scale bars correspond to 10 μm, and dashed lines indicate corresponding cross-sectional slices. **(D-F)** The distribution of **(D)** RI, **(E)** cell volume, and **(F)** dry mass of control, osmotically stressed, and osmotically adapted yeast cells, respectively. The box plot indicates the interquartile ranges (IQR) with a line at the median. The whiskers extend to the data within the 1.5 IQR of the respective quartile. Outliers outside of 1.5 IQR are marked as  (for each condition, *N* > 45).

Figure 3D shows that the mean RI value of yeast cells increases immediately upon osmotic stress, from 1.384±0.0002 (mean±SEM) to 1.419±0.0004, however, it recovers to 1.395±0.0003 as the cells adapt to osmotic pressure overnight. The RI changes are associated with the change of cell volume, as the yeast cells shrink upon osmotic stress from 65.77±1.91 fL to 44.34±1.66 fL and recover to a volume of 59.79±1.88 fL after the adaptation to osmotic pressure (Fig. 3E). Moreover, the dry mass of yeast cells increases upon osmotic stress from 14.98±0.38 pg to 18.21±0.69 pg and then maintains a similar dry mass of 17.69±0.55 pg during the osmotic adaptation (Fig. 3F). The initial increase of the dry mass after osmotic shock can be attributed to two processes. In short times the sorbitol molecule might diffuse into the cytoplasm, consecutively cells produce additional substances such as trehalose or glycerol as a protection mechanism from osmotic stress [29][30]. The addition of material leads to a slight increase of the mean RI value of osmotically adapted yeast cells compared to controls grown in YES medium. Here we show that cells adapt to and grow in a hypertonic medium despite high-density values even exceeding those exhibited by lowering intracellular pH. This finding implies that increasing intracellular mass density to values similar to, or even beyond those associated with lowering intracellular pH is not sufficient for keeping the cells in a growth-arrested state.

### Intracellular pH affects mass density recovery rate after osmotic stress

Our findings so far show that metabolic arrest induced by an acidified cytoplasm is correlated with higher intracellular mass density, while inducing the same high mass density by osmotic stress alone does not arrest cell growth. To confirm that only cells with low intracellular pH stay in the growth-arrested state and do not recover, we followed the recovery of osmotically stressed cells after changing their intracellular pH by measuring time-lapse RI tomograms. We induced osmotic stress on yeast cells by adding 1.2 M sorbitol to the phosphate buffer with 2% glucose in pH 7.5 (osmolality 2,029±32 mOsmol/kg) and pH 6.0 (osmolality 2,02±35 mOsmol/kg) as before, and then added DNP to equilibrate the intracellular pH to that of the surrounding medium. As a control, we added 1.2 M sorbitol to the YES medium. We then measured tomograms every 10 minutes in order to explore the change of mass density and volume of yeast cells after osmotic stress.

Figure 4A shows time-lapse tomograms of yeast cells with different intracellular pH when osmotic stress was induced at *t* = 0 minute. The control yeast cells (first row in Fig. 4A) recovered their RI value promptly after the osmotic stress, exhibiting an exponential decay of mean RI value. Meanwhile, the mean RI values of yeast cells with an intracellular pH of 7.5 (second row in Fig. 4A) stayed stable for the first 30 minutes (most probably for adapting to their transfer from YES medium into phosphate buffer medium), before the RI started to recover. More interestingly, the mean RI values of yeast cells with the intracellular pH of 6.0 (third row in Fig. 4A) did not change during the 60 minutes of observation (yellow box plots in Fig. 4B). While only increasing intracellular mass density at neutral pH was followed by a recovery of intracellular RI and mass density, the lack of such recovery in low intracellular pH implies that physical and mechanical changes of the cytoplasm, associated with growth arrest, have occurred.

**Figure 4.**
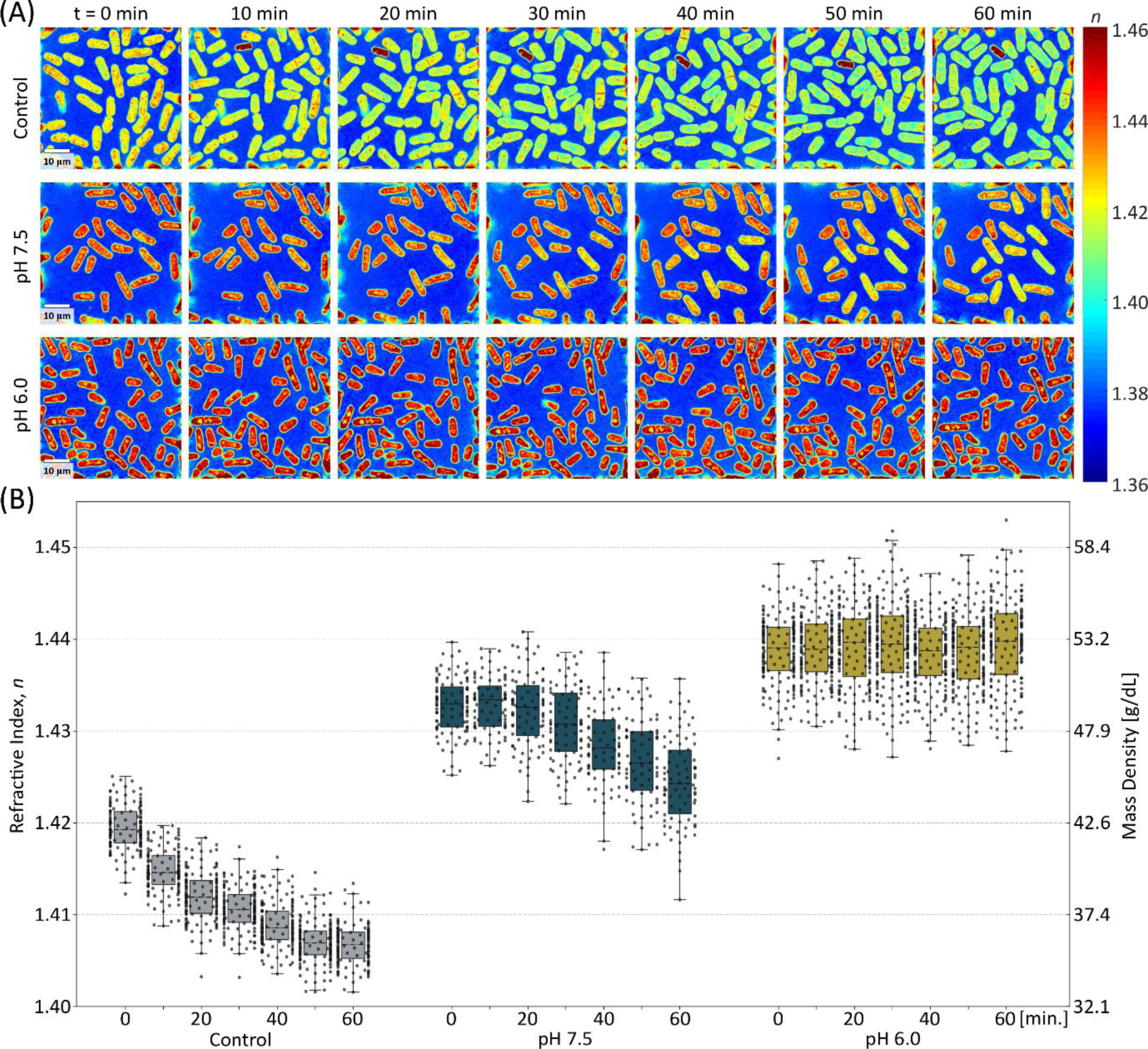
The effect of intracellular pH on the recovery rate of the mass density of yeast cells after osmotic stress. **(A)** The cross-sectional slices of time-lapse 3D RI tomograms of yeast cells with different intracellular pH, acquired every 10 minutes after osmotic stress. The scale bar indicates 10 μm. **(B)** The mean refractive index distribution of osmotically stressed cells in YES medium (gray, *N* > 120), at pH 7.5 (blue, *N* > 70) and at pH 6.0 (yellow, *N* > 150). The box plot indicates the interquartile ranges (IQR) with a line at the median. The whiskers extend to the data within the 1.5 IQR of the respective quartile. Outliers outside of 1.5 IQR are marked as ×.

To test the mechanical status of the fission yeast cytoplasm it is possible to remove the rigid cell wall (spheroplasting) and then to follow their rounding up - or the lack thereof [11]. As shown in Figs. 5(A-B), osmotically stressed yeast cells in YES medium and with a intracellular pH of 7.5 rounded up into a spherical shape to minimize the surface to volume ratio after spheroplasting, which indicated that the cytoplasm was in a liquid-like state [31,32]. They also had slightly lower RI values than osmotically stressed yeast cells without spheroplasting, which might have induced an increase in volume. However, osmotically stressed yeast cells with an intracellular pH of 6.0 did not acquire a round shape upon cell wall removal and exhibited increased RI values, which demonstrated the transition of the cytoplasm into a solid state with an inherent elastic stiffness that prevented rounding up.

**Figure 5.**
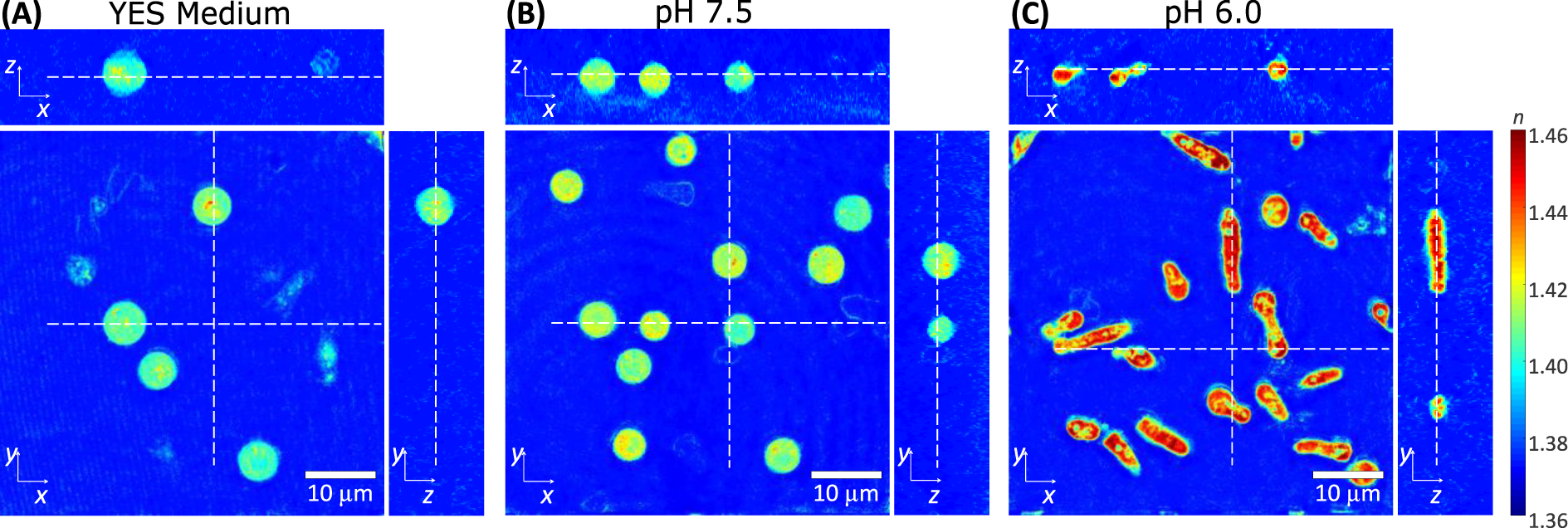
The cross-sectional slices of refractive index tomograms of yeast in **(A)** YES medium with 1.2 M sorbitol, **(B)** phosphate buffer in pH of 7.5 and **(C)** pH 6.0 with 1.2M sorbitol and DNP, after the removal of cell wall. The scale bar indicates 10 μm.

### Increased intracellular mass density is associated with but not sufficient for cell stiffening

We had previously demonstrated that the acidified cytoplasm undergoes a liquid-to-solid transition [11] and showed now that this coincides with an increase in the intracellular mass density. However, our data suggest that a sudden mass density increase of the cytoplasm may not be sufficient to promote this transition. In order to quantify and relate the mechanical changes to mass density changes in the yeast cytoplasm, we carried out AFM indentation experiments and ODT measurements of osmotically stressed and adapted yeast cells taken from identical batches. To extract and compare the mechanical properties of yeast cytoplasm using a Hertz model with AFM, it is important to apply the method to the same spherical geometry. This can be achieved by first removing the cell wall of *S. pombe* and then changing the intracellular pH. However, removal of the cell wall also requires the use of high molalities of sorbitol to prevent cell lysis.

Thus, to measure the effect of intracellular pH only, it is important to adapt the cells to these high molarities. To this end, we first measured spheroplasts of cells that had been adapted to hypertonic medium overnight before cell wall removal and adjustment of pH, as shown in Fig. 6A. Yeast spheroplasts with low intracellular pH had a higher RI, which is directly correlated with higher mass density, than with neutral pH as shown in Fig. 6B. Importantly, the apparent Young’s modulus of the acidified spheroplasts was significantly higher (*E* = 17.6±0.2 kPa; mean±SEM) compared to the Young´s modulus of spheroplasts with a neutral intracellular pH (*E* = 4.3±0.1 kPa) (Fig. 6C). These results demonstrate that acidification of the yeast cytoplasm coincides with mass density increase and an overall stiffness increase.

**Figure 6.**
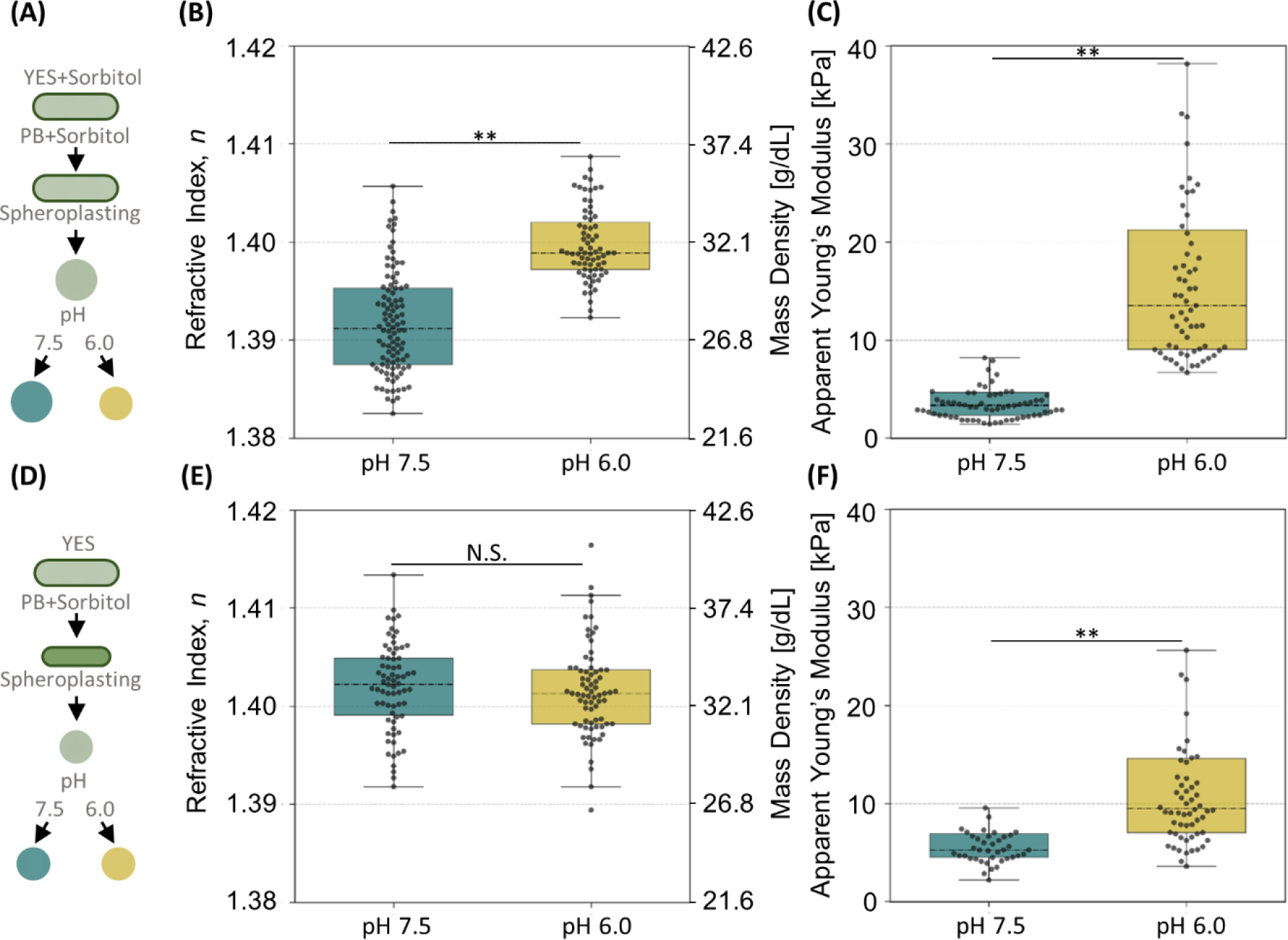
Stiffness and mass density of yeast cells with different intracellular pH during or after adaptation to osmotic stress. **(A, D)** Schematic diagram for cell culture, spheroplasting in phosphate buffer (PB) with sorbitol and pH adjustment for **(A)** overnight osmotically adapted and **(D)** osmotically stressed *S. pombe*. **(B, E)** The distribution of mean refractive index of yeast cells with intracellular pH of 7.5 and 6.0 in **(B)** osmotic adaptation (*N* > 70) and **(E)** osmotic stress (*N* > 70). **(C, F)** The distribution of the apparent Young’s modulus of spheroplasts in the intracellular pH of 7.5 and 6.0 in **(C)** osmotic adaptation (*N* > 60) and **(F)** osmotic stress (*N* > 40). The box plot indicates the interquartile ranges (IQR) with a line at the median. The whiskers extend to the data within the 1.5 IQR of the respective quartile. Outliers outside of 1.5 IQR are not shown.

To determine if an increased intracellular mass density is sufficient to stiffen the yeast, we osmotically stressed cells cultured in YES medium by addition of 1.2 M sorbitol, then immediately removed the cell wall, and subjected the spheroplasts to ODT and AFM indentation measurements (Fig. 6D). In agreement with the notion that sorbitol induces osmotic stress, the RI values of cells with a neutral intracellular pH already exhibited a high refractive index. Interestingly, the intracellular mass density of yeast spheroplasts with a low intracellular pH was not further increased (Fig. 6E). Importantly, the apparent Young’s modulus of the acidified spheroplasts was significantly higher (*E* = 13.1±0.2 kPa; mean±SEM) compared to the Young’s modulus determined for spheroplasts with a neutral intracellular pH (*E* = 6.3±0.1 kPa), despite the similar mass density (Fig. 6F). These results unambiguously demonstrate that increased intracellular mass density does not by itself lead to cell stiffening. Apparently, the cytoplasm requires, in addition to increased content crowding, other mechanisms to acquire its mechanical stability and to arrest growth.

## Discussion

In this study, we investigated the relationship between intracellular mass density, changes in intracellular pH and stiffness of growing and arrested *S. pombe* yeast cells. We adopted the method of Munder et al. for lowering intracellular pH in yeast cells [11]. We employed ODT to measure the RI values and evaluate the mass density of yeast cells, and found that mass density is increased in the acidified yeast cytoplasm. We then showed that increasing mass density triggered by osmotic stress is not sufficient to promote a permanent transition into a mechanically stiff and growth-arrested state. We confirmed that yeast cells with acidified cytoplasm exhibited characteristics of dormancy as they did not exhibit any distinct mass density change after osmotic stress and showed no rounding up upon cell wall removal. We assessed the mechanical properties of acidified and neutral yeast cytoplasm using AFM. We demonstrated that an increase in stiffness is accompanied with an increase in the intracellular mass density. However, increasing mass density of active yeast cells by osmotic stress was not associated with an increase in stiffness.

In this work, we have, for the first time, employed ODT to evaluate mass density changes and total dry mass of the yeast cytoplasm. We provided evidence for the mechanism of yeast cells to lose water and increase mass density when dormancy is induced by lowering intracellular pH. This crowded intracellular content leads definitely to a change in the physical properties of the cytoplasm that were so far explored using traditional techniques such as exogeneous particle tracking [11,33,34]. Various explanations have previously been proposed for the nature of these changes. Some suggested that the cytoplasm undergoes a glass transition due to increased mass density [35]. Others described a cytoplasm as water-containing matrix formed by a cytoskeletal network that is behaving as a hydrogel [36]. We here demonstrate that increasing the mass density of cells will not by itself result in stiffening and growth arrest of yeast cells. We show that the cytoplasm of metabolically arrested cells acquires, along with an increased mass density, solid-like characteristics. We have also explored that different degrees of compression increased the crowding of the cytoplasm using particle tracking method (see Supplementary Figure 2). We thus suggest that the combination of high mass density and reinforcing the cytoplasm with some percolated network of macromolecular assemblies is necessary for providing the cells with the mechanical stability needed for enduring harsh environmental conditions and ultimately survival.

Still, in order to profoundly unravel the structure and the organization of the solid cytoplasmic network, it is necessary to explore the response of these physical properties at various time scales. This could be achievable with the use of AFM dynamic probing of the mechanical properties at different frequencies. Using AFM provides a straight-forward measurement of cytoplasm, nevertheless, it requires steps of removing the yeast cell wall, which could potentially alter cell properties, and current analysis assumes homogeneity of the sample. Thus, there is a growing interest in using noninvasive optical methods to probe mechanical properties of soft materials such as Brillouin microscopy [37,38] and the temporal correlations of time-lapse quantitative phase microscopy [39,40].

Taken together, we present here a study that augments the field of phase transitions in dormant organisms with an insight into the optical and mechanical properties of the cytoplasm. We defined the interplay between intracellular mass density, stiffness and dormancy. Further studies for exploring the key factors and the organization of the solid cytoplasm will be essential to understand the underlying mechanisms of these organisms to initiate dormancy.

## Author Contributions

SAb, KK, TF, AE, RS, SM, HK, SAl, VZ, JG contributed to the conception and design of the study; SAb prepared the cells, conducted the AFM measurements, analyzed and interpreted AFM data; KK prepared the cells, conducted the ODT measurements, analyzed and interpreted ODT data; TF, AE provided the cells and the mediums, provided biological support for experiments; DM performed particle tracking, DM and VZ interpreted particle tracking data; RS and SM contributed to experiment performance; HK and VZ contributed to the theoretical aspect of data interpretation; SAb and KK wrote the first draft of the manuscript; TF, DM, VZ and JG contributed to manuscript revision.

## Acknowledgments

We thank the Microstructure Facility at the Center for Molecular and Cellular Bioengineering (CMCB) at Technische Universität Dresden and JPK Instruments AG in Berlin for excellent support. We are also grateful to Gheorge Cojoc, Joan-Carles Escolano, Paul Müller, Anna Taubenberger, Torsten Müller, Vamshidhar Gade, Teymuras Kurzchalia, Elisabeth Fischer-Friedrich and Kamran Hosseini for helpful discussions and technical support.

## Data Availability Statement

Datasets are available on request

